# Soft tissue morphology of the vomeronasal organ in *Lontra canadensis* and its osteological correlate: Implications for the evolution of the caniform accessory olfactory system

**DOI:** 10.64898/2026.01.19.700381

**Authors:** Stephanie M. Palmer, William Foster, Grace Capshaw, Margot Michaud, Siobhán B. Cooke

## Abstract

The olfactory system plays a critical role in mammalian environmental perception, with some clades relying on an expanded accessory olfactory (vomeronasal) system (VNS) to detect larger, non-volatile odorants. Mammals make extensive use of this system for social communication between conspecifics. Recent studies have begun to investigate how the VNS changes in response to or as part of ecological transitions. Several studies have identified trends of VNS-associated gene loss or regression in secondarily aquatic mammals. However, continuing discussion on genotype-phenotype correlation within the VNS means that greater effort should be made to investigate the morphology of the VNS in species where it remains poorly understood. Here, we use skeletal and soft-tissue data to demonstrate that the vomeronasal groove, an established osteological correlate for the VNO in bats and primates, is also a valid indicator for its presence in Caniformia. Additionally, we confirm the presence of the VNO in the secondarily aquatic North American river otter (*Lontra canadensis*) and compare its morphology with that of two close-related species, the semi-aquatic American mink (*Neogale vison*) and the terrestrial long-tailed weasel (*Neogale frenata*). This study expands the valid taxonomic scope of the vomeronasal groove’s proxy as an osteological correlate, confirms the presence of the VNO in the previously undescribed system of the North American river otter, and highlights the complexity of the mammalian accessory olfactory system.

## Introduction – Background

Vertebrate sensation is supported by a suite of specialized organs that allow organisms to perceive and process environmental stimuli. A key component of this sensory repertoire is the olfactory system, which detects odor molecules in the environment and communicates information to the forebrain for processing. Olfaction is functionally relevant to numerous fundamental vertebrate behaviors, including navigation (Poo et al., 2022), prey tracking (Hughes et al., 2010), reproduction (Aron, 1979; Baum & Kelliher, 2009; Baum & Cherry, 2015). The mammalian olfactory system is highly variable across species, reflecting anatomical and functional constraints (e.g., limitations related to skull morphology and/or respiratory function) as well as adaptation to diverse ecological niches. Understanding the evolutionary response (or contribution) of crucial sensory systems during major ecological transitions is a major goal of comparative biology and demands thorough sampling of phylogenetically relevant phenotypes, especially in poorly known taxa.

In mammals (and tetrapods generally), olfaction is functionally and anatomically segregated into the main olfactory system (MOS) and the vomeronasal system (VNS), also referred to as the accessory olfactory system (AOS) (Figure 1) (but see Baum, 2012 and Baum & Kelliher, 2009 for potential for functional overlap between the two systems). The special sensory function of the MOS is supported by the olfactory epithelia of the nasal cavity, which contain olfactory sensory neurons whose axons form the first cranial nerve (CN I) and are connected directly to the olfactory lobes of the brain. The anatomy of the VNS, in contrast, is much more complex.

**Figure 1.**
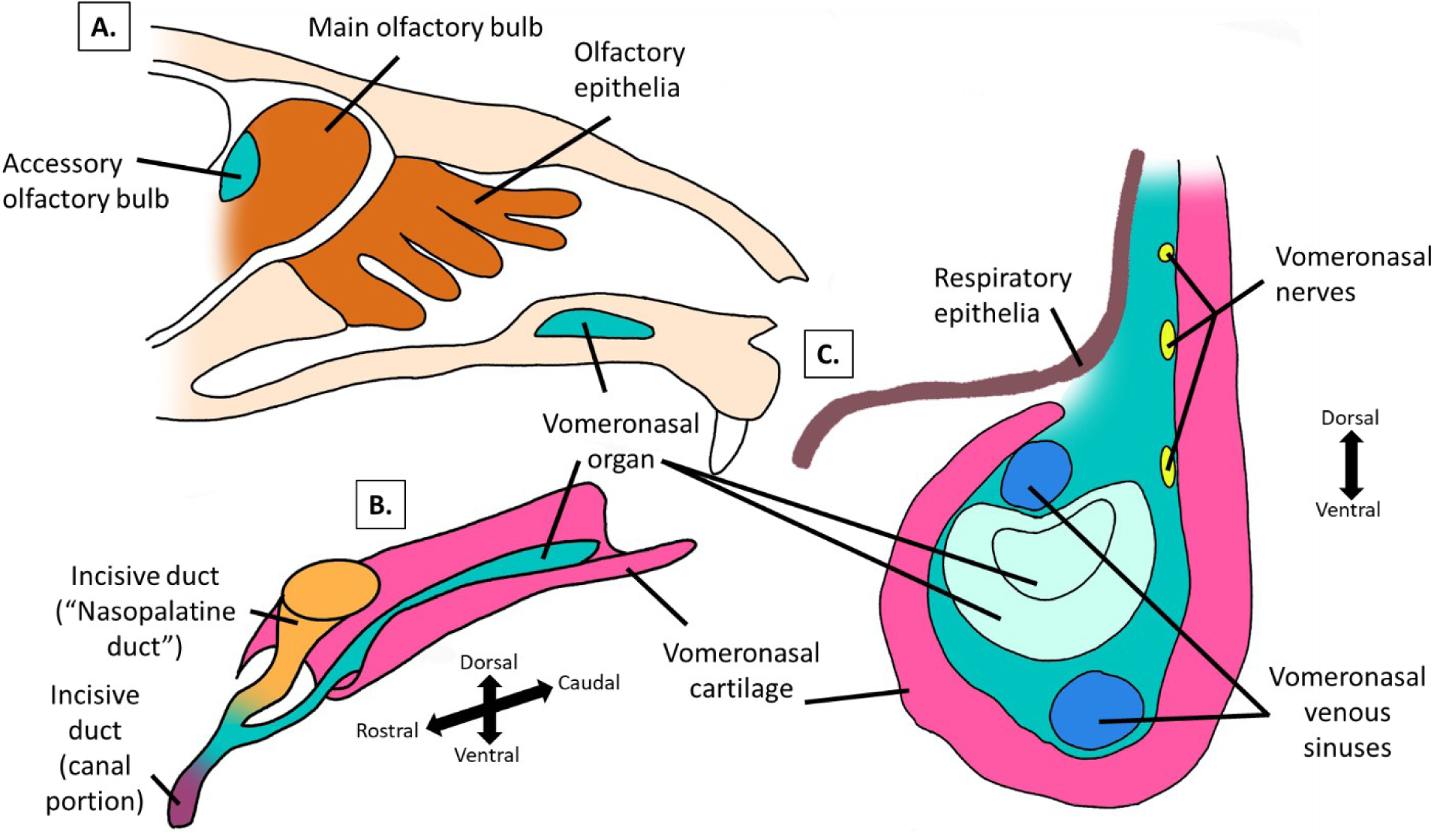
Drawings of the gross anatomy in parasagittal view (A), sub-gross anterolateral view (B), and coronal cross-sectional view (C) of the histo-morphological anatomy of the main olfactory system (MOS), accessory olfactory system (AOS), and vomeronasal organ (VNO). **A**: the main olfactory bulb and olfactory epithelia represent the MOS (brown-orange), while the vomeronasal organ (VNO) and accessory olfactory bulb represent the AOS (blue-green). **B:** relationship of the VNO, vomeronasal cartilage (VNC), and incisive (nasopalatine) duct in a sub-gross anterior-lateral orientation, with directional axes indicated (dorsal–ventral, rostral–caudal). **C:** coronal cross-section of the VNO, highlighting the vomeronasal sensory epithelium, respiratory epithelium, vomeronasal nerves, and vomeronasal venous sinus with directional axes indicated (dorsal-ventral). Modified after Salazar and Sánchez-Quinteiro (2009), Evans (1993), and Estes (1978).

The VNS is supported by specialized epithelia that are organized into a pair of parasagittal organs located lateral to the base of the nasal septum on the maxilla: the vomeronasal organs (or Jacobson’s organs) (VNO) (Salazar et al., 2003; Garrett, 2015; Smith et al., 2024). The VNO is supported by a cartilaginous, sometimes ossified, capsule, referred to as the vomeronasal cartilage (VNC), which protects and provides structural support to the organ (Salazar et al., 1995). The VNO capsule communicates anteriorly with the incisive duct (ID), which travels through the incisive canal and opens into the oral cavity (Figure 1; Ortiz-Leal et al., 2024). The VNO is innervated by vomeronasal nerves (VNN) that originate from functionally distinct receptor cells from the MOS. The VNN connects the VNO to a specialized region of the olfactory lobes, the accessory olfactory bulbs (AOB). Whilst the MOS is responsible for processing small, airborne, and molecularly simple odorants, the VNS has been linked to the processing of larger, non-volatile odorants, such as social chemosignals (“pheromones”) (Garrett, 2015), and is thus hypothesized to be critical for communication between conspecifics, particularly concerning individual recognition and sexual reproduction (Zufall et al., 2002; Keverne, 2004; Tirindelli, 2021).

Evidence for reductions of the MOS and AOS-related homologues in secondarily aquatic lineages often support the hypothesis that land-to-water transitions universally result in reduced olfactory acuity (Yu et al., 2010; Berta et al., 2014; Lu et al., 2016; Paulina-Carabajal et al., 2015; Zhang & Nikaido, 2020) and may be compensated for by an expansion of other sensory systems, e.g., the “visual-priority hypothesis” (Debey & Pyenson, 2013; Garrett, 2015; Hadden & Zhang, 2023). One of the most studied aspects of the AOS is the underlying genetic components, in which a reduction in AOS-related genes, including VNO receptor genes (ancV1R; TRPC2), has been identified across multiple lineages of secondarily aquatic mammals (e.g., Lutrinae, Pinnipedia, Sirenia, Cetacea; Yu et al., 2010; Hecker et al., 2019; Liu et al., 2019; Zhang & Nikaido, 2020) and bats (Zhao et al., 2011).

Other anatomical and genetic transitions in the MOS and VNS in secondarily aquatic mammals have also been identified (Mackey-Sim et al., 1985; Pihlström et al., 2008; Berta et al., 2014; De Vreese, 2023, Farnkopt et al., 2025), including several within secondarily aquatic Carnivora (Switzer et al., 1980; Zhang & Nikaido, 2020 Kondoh et al., 2024; 2025). Pinnipeds (seals, sea lions and fur seals, and walruses) have derived morphologies in the VNS (Switzer et al., 1980; Kondoh et al., 2024; 2025), mirroring a reduction in VNO receptor genes (Yu et al., 2010; Zhang & Nikaido, 2020). Studies of VNS anatomy in secondarily aquatic carnivorans (Table 1) have revealed the absence of an AOB in the true seal species *Phoca vitulina* (Switzer et al., 1980) along with the modification of its VNO from a sensory to a secretory structure (Kondoh et al., 2024; Figure 2B). Meanwhile, members of Otariidae, including *Callorhinus ursinus, Eumatropias jubatus,* and *Zalophus californianus*, retain an AOB (Switzer et al., 1980), with Steller sea lions (*E. jubatus*) retaining a functional VNO (Kondoh et al., 2025; Figure 2C). The Steller sea lion VNO is derived from the ancestral condition through its migration from the ventral surface of the nasal cavity to within the incisive canal (Kondoh et al., 2025).

**Figure 2.**
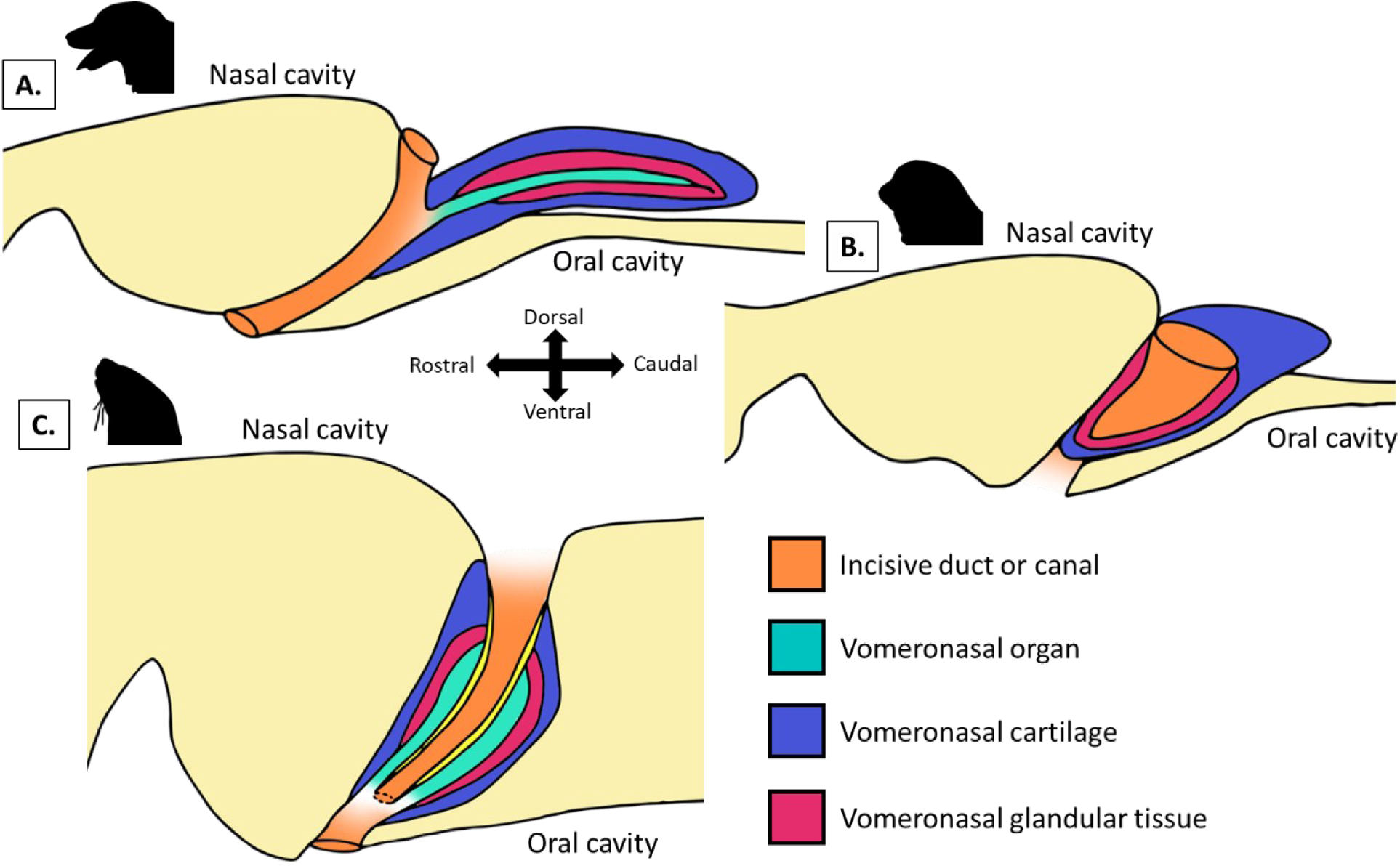
Comparative parasagittal illustrations showing shows the spatial relationships between the incisive duct or canal (orange), vomeronasal organ (cyan), vomeronasal cartilage (blue), and vomeronasal glandular tissue (magenta) within the nasal and oral cavities. Cranial bones colored in yellow. Panel A: a terrestrial carnivoran (*Canis familiaris*), Panel B: a phocid (Harbor seal, *Phoca vitulina*), and Panel C: an otariid (Steller sea lion, *Eumatopias jubatus*). Directional axes (rostral-caudal and dorsal-ventral) are indicated in panel A. Adapted from Kondoh et al., 2024 and 2025. Silhouettes from PhyloPic (all dedicated to the public domain): *Phoca vitulina, Eumatopias jabatus* by Andy Wilson; *Canis familiaris* by Tracy Heath.

**Table 1.**
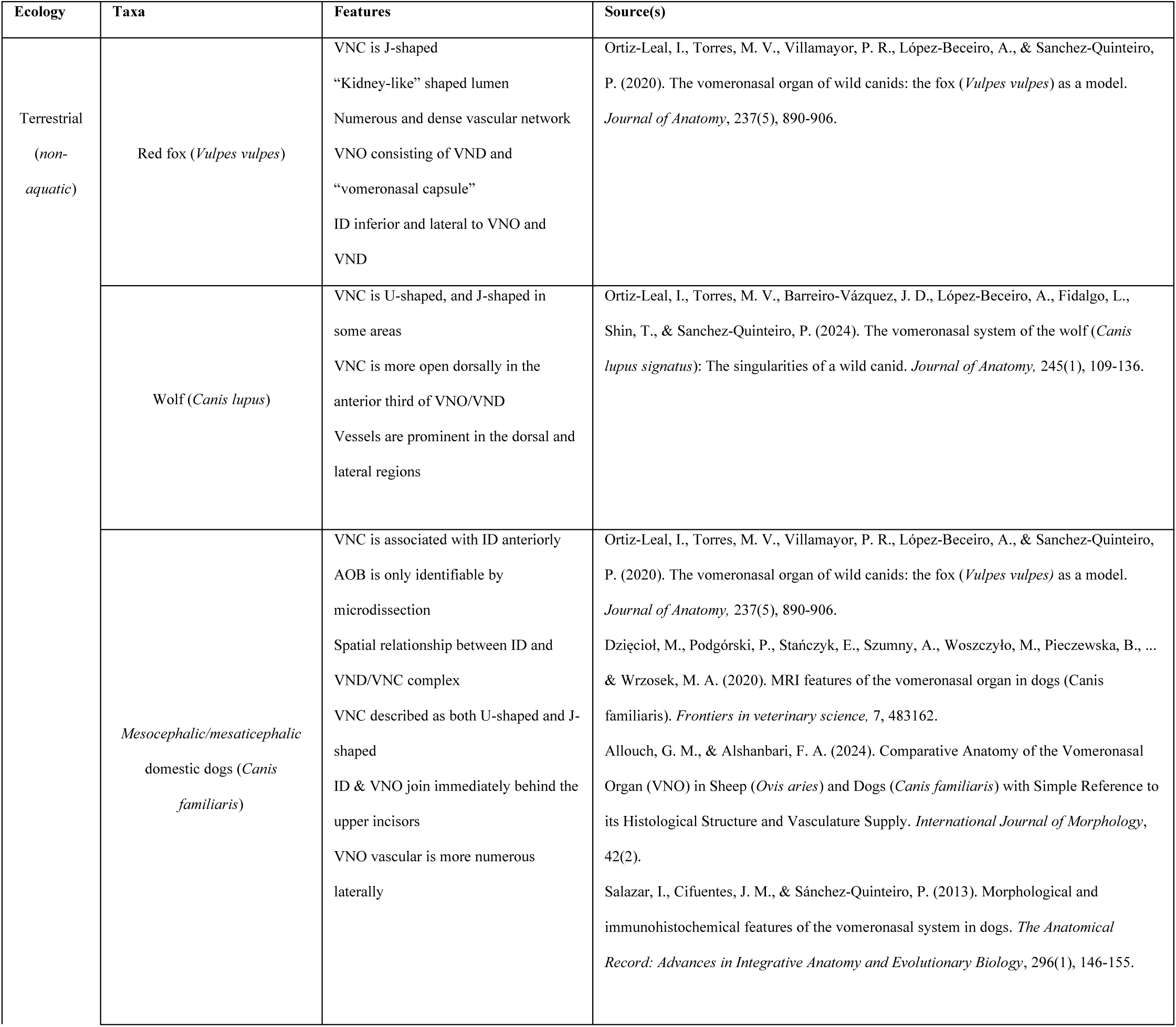
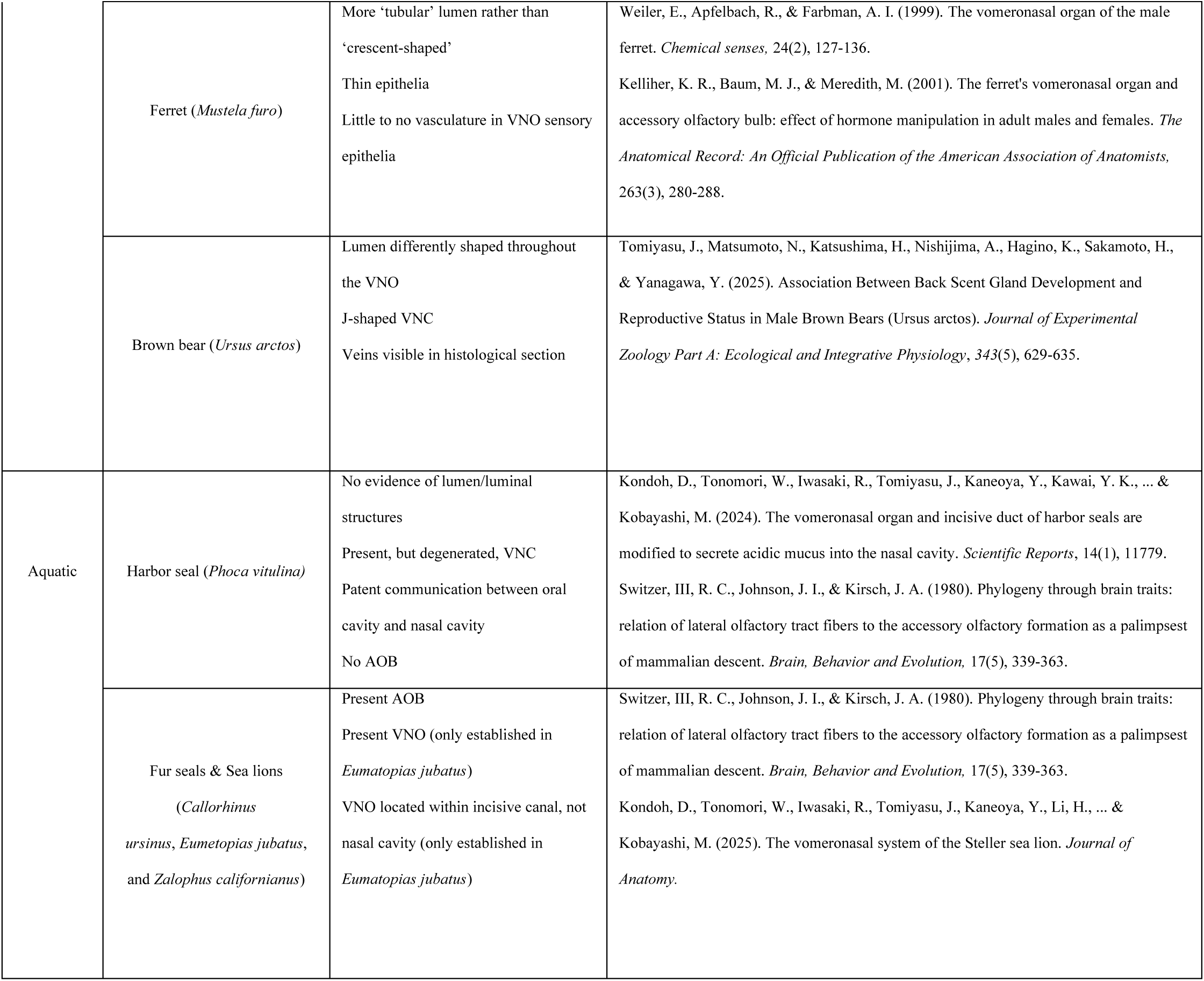
Major relevant comparative features of AOS structures across Caniformia, including pinnipeds. VNC = vomeronasal cartilage; VNO = vomeronasal organ; VND = vomeronasal duct; ID = incisive duct; AOB = accessory olfactory bulb.

The other secondarily aquatic caniforms, otters (Lutrinae), possess reduced VNO receptor gene repertoires similar to the condition observed in phocid pinnipeds - specifically sea otters (*Enhydra lutris*), giant river otters (*Pteronura brasiliensis*), and the Eurasian otter (*Lutra lutra*) (Yu et al., 2010; Hecker et al., 2019; Zhang & Nikaido, 2020). Some authors hypothesize that it is likely that lutrines do not possess a VNS or VNO, similar to phocid pinnipeds (Zhang & Nikaido, 2020); however, no information has been presented on whether their anatomical components converge on a similar derived morphological condition (Yu et al., 2010; Hecker et al., 2019; Zhang & Nikaido, 2020; Zellmer et al., 2021). While some studies have found a strong correlation between genetics and morphology within the MOS and VNS (Garrett & Steiper, 2014; Bird et al., 2018; Farnkopt et al., 2025), other studies show that phenotypes exhibit stability despite reductions in receptor gene repertoires (Niimura & Nei, 2007; Yohe et al., 2020; Yohe & Krell, 2023). As a result, conclusions on the strength and nature of correlations between olfactory genes and phenotype vary across studies (Niimura & Nei, 2007; Salazar & Sánchez-Quinteiro, 2009; Yohe et al., 2018; Yohe et al., 2020; Yohe & Krell, 2023), and the nature of phenotype-genotype correlation is often system- and scale-dependent (Garrett & Steiper, 2014; Yohe et al., 2020). The emerging complexity of the correlation between genotype and phenotype in this system is not only intellectually motivating in its own right but also touches on broader themes of phenogenetic drift (Weiss & Fullerton, 2000) and mosaicism (Caianiello, 2024). Ultimately, studying VNS morphology in species for which knowledge gaps exist is critical to understanding the relationship between phenotype and genotype in the mammalian olfactory system.

To further evaluate the phenotype of the Lutrine VNS, we utilize both soft-tissue and osteological data. To examine soft-tissue, we used diffusible iodine contrast-enhanced micro-computed tomography (diceCT) scans. DiceCT is a method that uses contrast agents (Lugol’s iodine and/or Strong solution) to visualize soft tissues in situ, with adjacent and underlying hard tissues (e.g., bone), in CT data (Gignac et al., 2016). Its application in visualizing the soft tissues of the VNO has been validated in recent publications, with authors noting its potential for identifying small structures such as the VNO (Yohe et al., 2018; Smith et al., 2021; Smith et al., 2024; Collin et al., 2024; Leng & Shi, 2025). Our diceCT dataset capturing soft-tissue variation of the VNO around Lutrinae consisted of three mustelid species: American mink (*Neogale vison*), least-tailed weasel (*Neogale frenata*), and North American river otter (*Lontra canadensis*). To contextualize the morphology observed, we also consulted the literature to define common features of the VNO across caniforms (Table 1). While it is important to mention that genes associated with the VNS have not yet been explicitly characterized in the North American river otter, phylogenetic bracketing (Witmer & Thomason, 1995) supports the inference of similar genetic reductions based on its relationship to the studied taxa *Enhydra lutris*, *Pteronura brasiliensis*, and *Lutra lutra* (Waku et al., 2016; de Ferran, 2022).

In addition to soft-tissue morphology from diceCT, other methods assess the presence and morphology of the VNO by investigating its osteological correlate, the vomeronasal groove (VNG). Morphologically, the VNG is described as a bilateral, rostral-caudally elongate, and medial-laterally narrow trough or indentation located on the nasal septum, with a raised lateral edge and an impressed center (Smith et al., 2011; Garrett et al., 2013; Garrett, 2015; Smith et al., 2024). Notably, previous work has established the presence of such structures in both extant and fossil mammals (Hillenius, 2000; Crompton et al., 2017; Bendel et al., 2018), as well as in detailed comparative studies establishing it as a formal correlate in primates (Smith et al., 2011; Garrett et al., 2013; Garrett, 2015) and bats (Smith et al., 2024). Both the presence and dimensions of the VNG have been shown to correlate with the presence or size of the VNO (Garrett, 2015), revealing evolutionary patterns of changes to the AOS, including diverse trajectories of degeneration seen in bats (Smith et al., 2024).

To date, no study has used VNG as an indicator of VNO presence in carnivorans, particularly in the application to understudied secondarily aquatic taxa such as Lutrinae. In reviewing existing histological and anatomical studies on the VNO morphology in canids (e.g., *Vulpes vulpes*, *Canis lupus*, and [mesocephalic] *Canis familiaris*), ursids (*Ursus arctos*), and mustelids (*Neogale vison*, *Mustela furo*) [Salazar et al., 1995; Weiler et al., 1999; Kelliher et al., 2001; Tomiyasu et al., 2017; Mahdy & Mohamed, 2019; Ortiz-Leal et al., 2020; Dzięcioł et al., 2020; Ortiz-Leal et al., 2022; Ortiz-Leal et al., 2024; Sanmartín-Vázquez et al., 2024; Table 1]), a groove-like indentation in the ventral nasal cavity, on either side of the nasal septum or vomer, was present and in close association with the VNO and VNC, suggesting it is homologous to the VNG described in primates and bats (Smith et al., 2011; Garrett et al., 2013; Garrett, 2015; Smith et al., 2024). This evidence from the literature suggests that the VNG is a strong candidate as an osteological correlate for the functional VNO in caniforms, expanding its utility for estimation of VNO presence in other otter taxa (e.g., *Lutra, Aonyx, Lutragale, Enhydra*) not represented by diceCT data.

## Methods

### Materials: Dry Skulls

This study is based on a comparative sample across Caniformia, selected to capture taxonomic and ecological diversity within the clade, as well as size variation. CT scans were sourced from MorphoSource and from scans taken by Arianna Harrington and Matthew Colbert using CT scanners at the Duke Shared Materials Instrumentation Facility (SMiF) (Duke University) and the University of Texas High-Resolution X-ray Computed Tomography Facility. Specimens were sourced from the National Museum of Natural History (MNHN), Smithsonian Museum of Natural History (USNM), Laboratory of Santiago Palazón (LSP), Yale Peabody Museum (YPM), Illinois State Museum (ISM), University of Florida Museum of Natural History (UF), Duke Evolutionary Anthropology Department (DU), Royal Belgian Institute of Natural Sciences (RBINS), The Pennsylvania State University Department of Anthropology (PSU), North Carolina Museum of Natural Sciences (NCSM), University of Alaska Museum (UAM), and the American Museum of Natural History (AMNH) (Supplemental Tables 1 and 2). This dataset includes specimens representing 45 species across fissiped caniforms. Most species were represented by a single specimen. The nasal and oral cavities of these CT-scanned specimens were examined using 3DSlicer (Federov et al., 2012) to identify evidence of VNGs in coronal and 3D-rendered views. CT scans were loaded into 3DSlicer using the ImageStacks module of the SlicerMorph package (Rolfe et al., 2021). Some scans were imported at reduced resolution or with the “skip slices” (1-2 slices) option to enhance loading performance. Figures were generated using the Capture feature in 3DSlicer, Microsoft PowerPoint, and Adobe Photoshop. Standard adjustments to contrast and brightness were made in these applications to enhance visibility for the identification of structures during analysis and figure preparation.

### Materials: Heads & Soft Tissue

All procedures for acquiring soft-tissue specimens and associated data adhered to ethical standards. The river otter (*Lontra canadensis*) and least weasel (*Neogale frenata)* diceCT scans were downloaded from MorphoSource (Supplemental Information Table 2). The American mink (*Neogale vison)* was sourced from Carolina Biological Supply as a whole-body, skinned specimen preserved in Carolina Perfect solution. Decapitation was achieved via atlantooccipital dislocation prior to CT scanning. Scans were generated using a microCT RX Solutions EasyTom scanner at the Johns Hopkins University Materials Characterization and Processing Center. The specimen was scanned at 48.5 µm voxel resolution before staining and re-scanned at 16.5 µm voxel resolution after 41 days in 3.75% buffered Lugol’s iodine solution (Gignac et al., 2016; Dawood et al., 2021). Scans were viewed in 3DSlicer, where structures were identified in coronal, sagittal, axial, and 3-dimensional views.

## Results

### Osteological Anatomy of the VNG - CT and Gross Morphology (Dry Skulls)

All species or specimens sampled exhibited evidence of a VNG. These structures were bilateral and consisted of rostro-caudally elongated and medio-laterally narrow grooves with indented centers and raised edges. VNGs were best visualized in coronal CT slices and 3D renderings of segmentations (Figures 3 and 4). The visibility of the VNG in 3D renderings varied depending on its position relative to surrounding anatomical structures, including the vomer (Figure 3). In some cases, the VNG could be visualized from a dorsal-anterior angle into the nasal cavity, although the vomer sometimes obstructed a direct view. When the VNG was not externally visible from dorsal-anterior views into the ventral nasal cavity floor, the nasal bones, frontal, or vomer were digitally cropped in an axial plane above the ventral floor of the nasal cavity, and/or the specimen was laterally rotated to visualize it. In addition to aligning with descriptions of the VNG in other mammals, remnants of VNC were visible in one specimen (*Mustela nivalis*, NCSM 8102; Figure 5).

**Figure 3.**
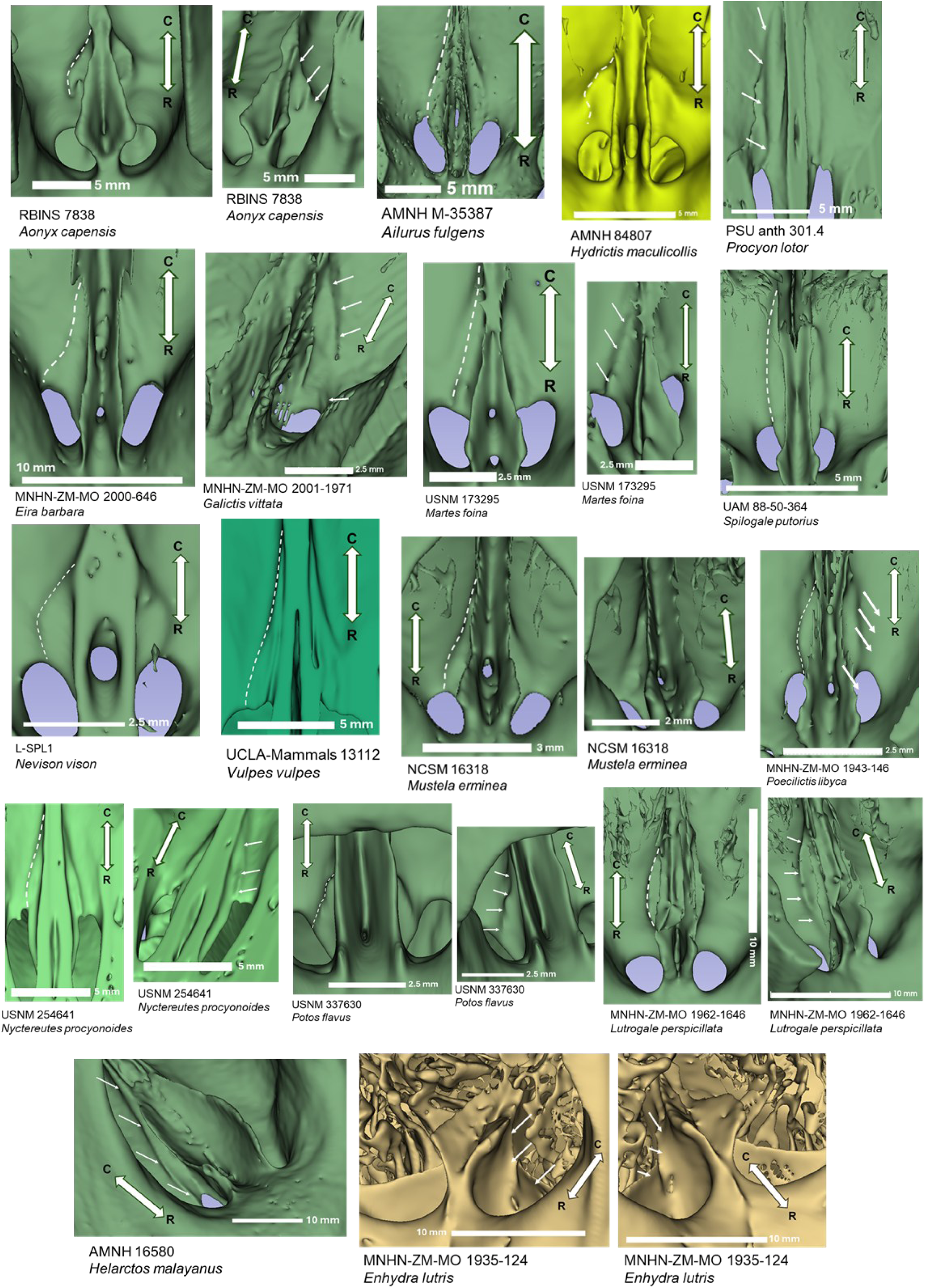
Dorsal and oblique views of the ventral surface of the nasal cavities from 3D renderings illustrating diversity in vomeronasal groove (VNG) shape; VNGs are traced in dashed lines on the right side in dorsal views, and oblique views show arrows pointing towards the lateral margin of the VNG. Directional axes (rostral [R]–caudal [C]) are indicated in each image.

**Figure 4.**
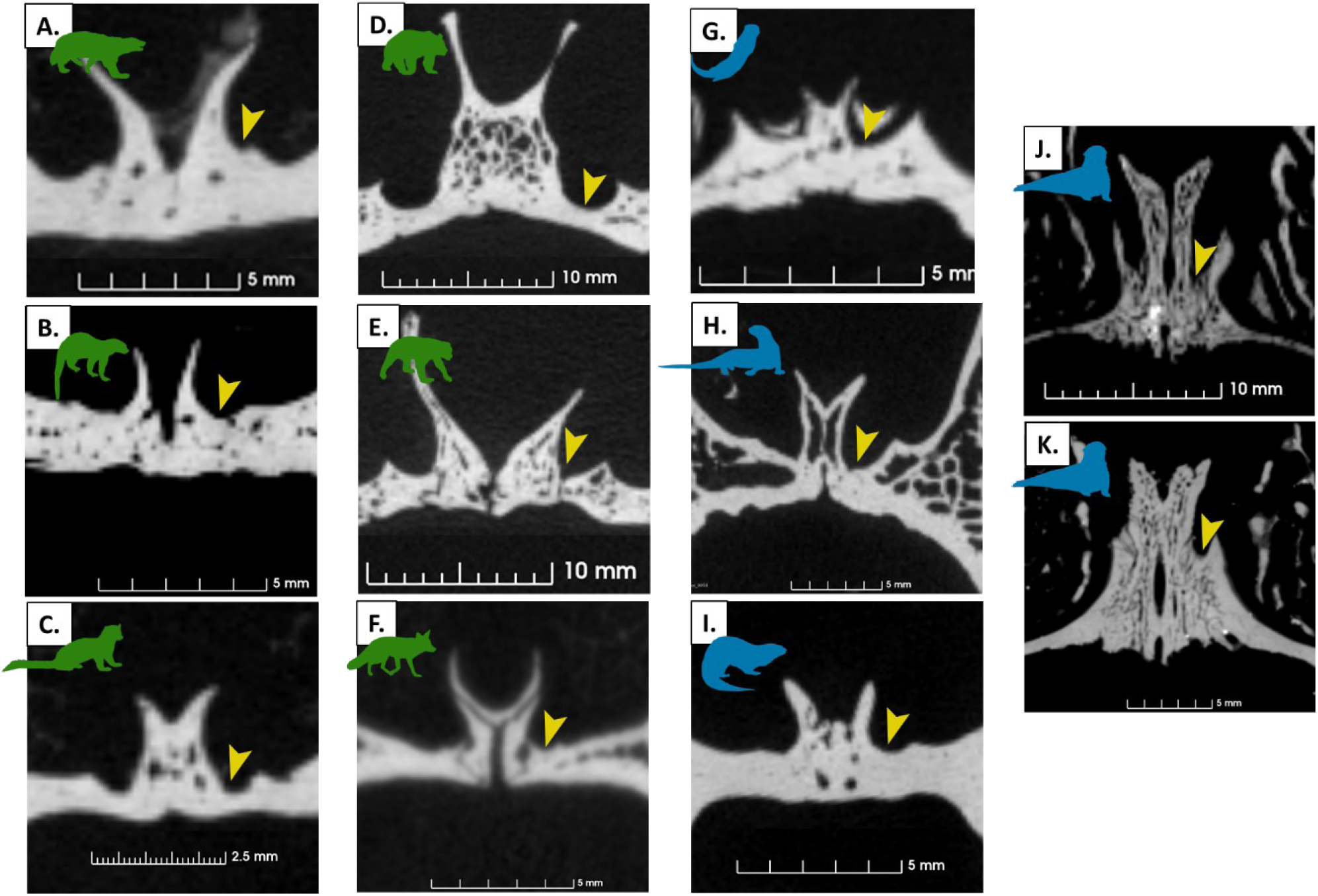
Coronal views of microCT scans showing the vomeronasal grooves across a diversity of caniform taxa. Color-coded silhouettes indicate ecological groups (*green* = terrestrial/arboreal taxa; *blue* = semi-aquatic taxa). Yellow arrowheads indicate the vomeronasal grooves (VNG). A. *Gulo gulo* (AMNH 37433); B. *Eira barbara* (MNHN-ZM-MO 2000-646); C. *Martes foina* (USNM 173295); D. *Ailuropoda melanoleuca* (AMNH 89030); E. *Helarctos malayanus* (AMNH 16580); F. *Vulpes vulpes* (UCLA 13112); G. *Lutra lutra* (USNM 259466); H. *Pteronura brasiliensis* (USNM 304663); I. *Lontra canadensis* (NCSM 19662); J. *Enhydra lutris* (MNHN ZM MO 1-35-124); K. *Enhydra lutris* (MNHN ZM MO 1962-1647).

**Figure 5.**
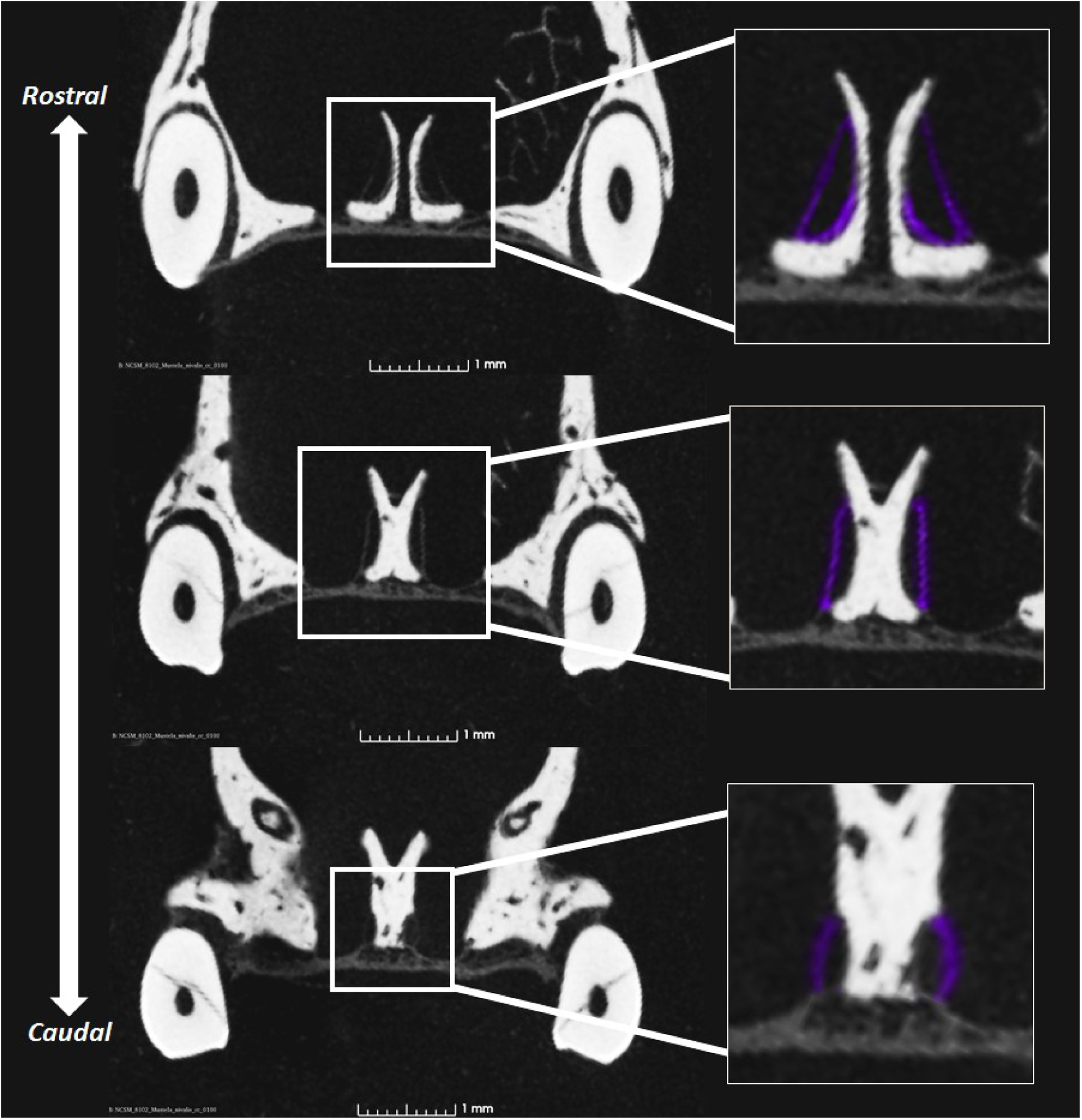
Three coronal CT slices of the nasal cavity of *Mustela nivalis* (NCSM 8102) showing remnant vomeronasal cartilage (VNC) in the dry skull. Slices progress from rostral to caudal from top to bottom. VNC is colored purple in the zoomed-in section.

The morphology of the grooves was variable and diverse across species, with most variation occurring in the groove’s cranial-caudal length, where some were longer and narrower (Figure 3G, E) and others were shorter and wider (Figure 3D, I). Longer VNGs tended to be mediolaterally narrower, while shorter or more abbreviated VNGs tended to be proportionally wider (Figure 3). The caudal end of the groove steadily (or in some cases, abruptly) narrowed towards the medial direction until its termination. In most species, the rostral termination of the VNG reached the incisive foramen, except for some Lutrine species (*Lutragale perspicillata*, Figure 3).

The VNG in all lutrines, except for the sea otter *Enhydra lutris*, largely resembled that of other mustelids but were notably shorter in the rostral-caudal dimension and oftentimes wider medio-laterally (Figure 3, Figure 4). The VNG of the semi-aquatic American mink (*Neogale vison*) was also rostral-caudally abbreviated (Figure 3). In one freshwater otter (*Lutragale perspicillata*), the VNG terminated early and did not reach the opening of the ID (Figure 3). In contrast, the VNG of the *E. lutris* exhibited a derived morphology relative to other species (Figure 3, Figure 4J and K). Specifically, it was dorsoventrally deep, mediolaterally narrow, and cranial-caudally shortened. The lateral margin was elevated, forming a distinct bony wall that bordered the groove. Unlike other caniforms, in which the lateral edge of the VNG is lower and merges gradually with the lateral portion of the palate or ventral nasal floor, the VNG in *E. lutris* was not continuous with these surrounding structures. Instead, the groove appeared more deeply inset, with a steeper lateral wall.

### diceCT Soft Tissue VNO Morphology - Mink, Long-tailed weasel, and Otter

In the diceCT scan of the mink (*Neogale vison*), the VNO is present and associated with a VNG structure (Figure 6). The VNO had a visible lumen that was dorsal-ventrally elongated and ovoid in cross-sectional shape. A patent incisive duct provided a connection between the oral and nasal cavities. Within the incisive duct, there was evidence of cartilaginous or mucosal tissue present at the periphery on the bony surface (a canal with a radiolucent center with radiodense tissue of differing densities along the walls) (Figure 6). In coronal, axial, and sagittal views, small circular radiolucent regions surrounding the VNO, distinct from the central vomeronasal duct (VND) lumen, were observed (Figure 6Cii; Figure 6Cv; white arrows). These structures were most numerous along the lateral and dorsal aspects of the VNO. They may represent either vascular or neural tissue, but are more likely vascular, as neural tissue typically appears radiodense in diceCT (Camilieri-Asch et al., 2020; Gignac et al., 2021). These features were present bilaterally and resemble the vascular VNO of *Vulpes vulpes* (Ortiz Leal et al., 2020). In the coronal view, the ID was ventrolateral to the VNO and VND. The lumen of the VND was generally ovoid across the entirety of its length, but some aspects of shape varied across the organ. Most variation in lumen shape was due to differences in the curvature of its superior and inferior margins, ranging from rounded to more angular forms. In the sagittal view, both the ID and the VNO were distinguishable from one another (Figure 6Cii).

**Figure 6.**
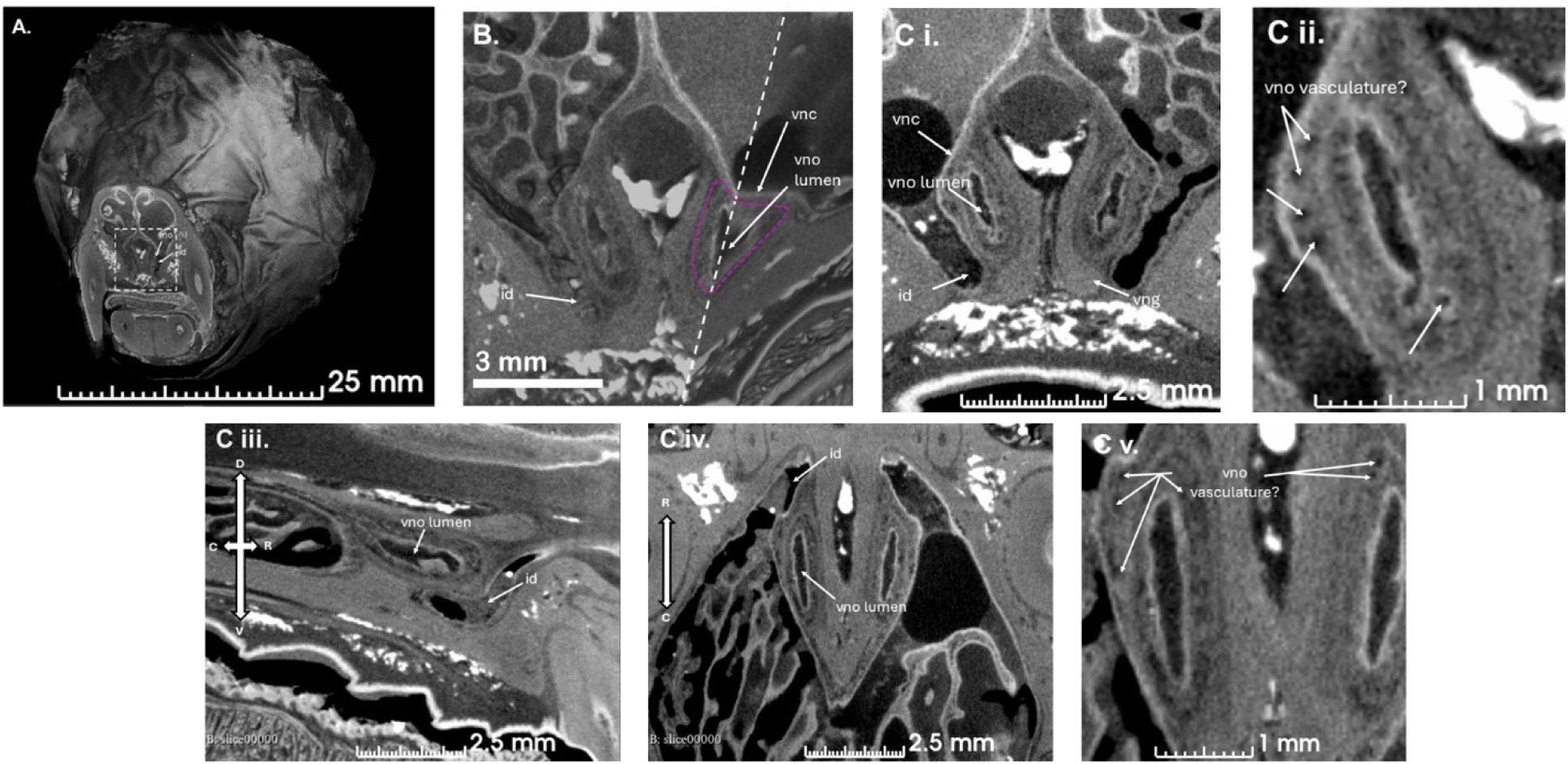
Views of diceCT of *Neogale vison,* magnified to see structures of the anterior nasal region and the vomeronasal organ. A. Full-head volume rendering. B. Volume rendering cropped to highlight the anterior nasal region in coronal and parasagittal view; the right vomeronasal organ is also cropped sagitally, indicated by the dashed line. C. Representative diceCT slices shown in **i, ii**. coronal, **iii**. parasagittal, and **iv, v**. axial planes. Directional axes are indicated in panels C iii and C iv. ID = Incisive duct; VNO = vomeronasal organ; VNC = vomeronasal cartilage.

Findings in the long-tailed weasel (*Neogale frenata*) were overall similar, with the presence of both a VNO, VNG, and ID (Figure 7). Differences in the *N. frenata* compared to *N. vison* include a larger, more dilated VND, VNO, and ID lumen, more distinct separation of the VND and ID lumen, and no visual evidence of VNO vasculature. The ID progresses from anterior to ventral-lateral relative to the VND and VNO, similar to *N. vison* (Figure 7).

**Figure 7.**
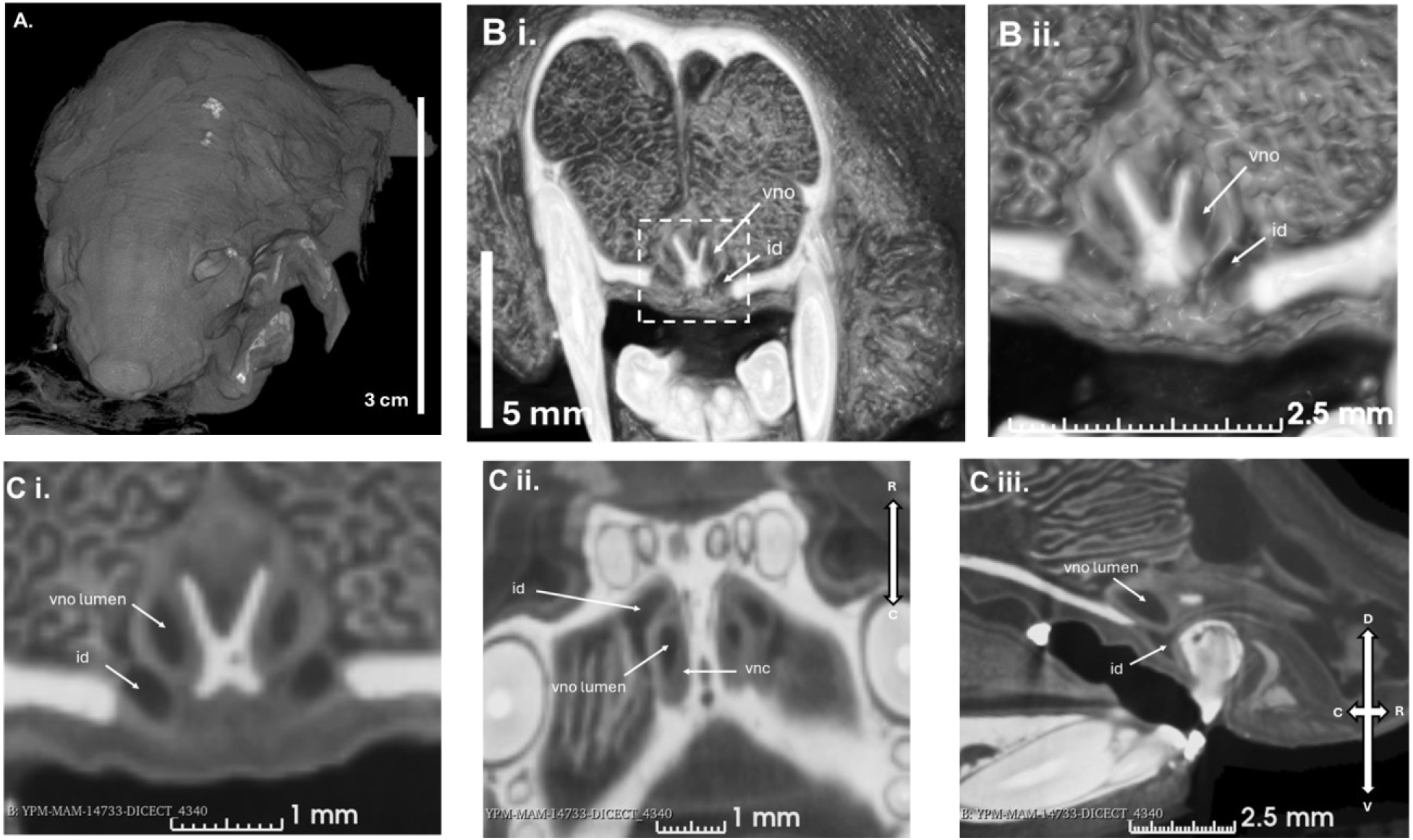
Views of diceCT of *Neogale frenata (YPM:VZ:YPM MAM 014733),* magnified to see structures of the anterior nasal region and the vomeronasal organ. A. Full-head volume rendering. B i. Three-dimensional volume rendering cropped in coronal section to highlight the anterior nasal region, and B ii. further magnified view showing additional detail. C. Representative diceCT slices shown in **i**. coronal, **ii**. axial and **iii**. parasagittal planes. Directional axes are indicated in panels C ii and C iii. ID = Incisive duct; VNO = vomeronasal organ; VNC = vomeronasal cartilage.

A VNO was present in the North American river otter (*Lontra canadensis)* (Figure 8). As in *N. frenata*, there was no visual evidence of VNO vasculature. An incisive duct (ID) was discernible from the VNO in multiple views and was generally similar to those of *N. vison* or *N. frenata* (Figure 8). In *L. canadensis*, the VND lumen was subtler, exhibiting lower contrast relative to surrounding tissue and being smaller in size compared to those of *N. vison* and *N. frenata*. Additionally, the VNO in *L. canadensis* was surrounded by more open space between it, the surrounding capsule, and the VNG than in either *N. vison* or *N. frenata*.

**Figure 8.**
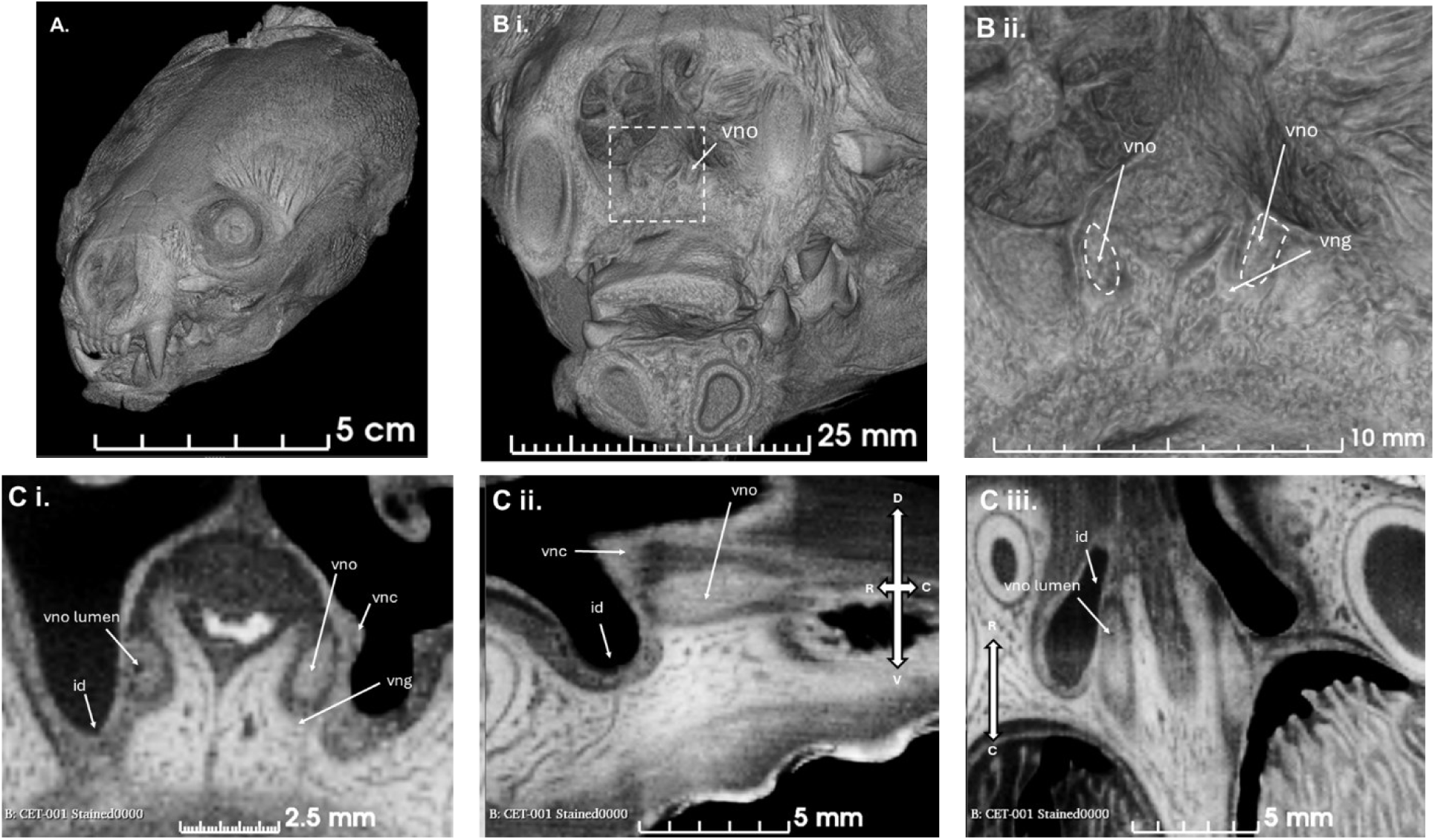
Series of coronal slices of diceCT of *Lontra canadensis (l-cet:001),* magnified to see structures of the anterior nasal region and the VNO. A. Full-head volume rendering. B i. Three-dimensional volume rendering cropped in coronal section to highlight the anterior nasal region, and B ii. further magnified view showing additional detail with VNO traced in white dashed lines bilaterally. C. Representative diceCT slices in i. coronal, ii. parasagittal, and iii. axial planes. Directional axes are indicated in panels C ii and C iii. ID = Incisive duct; VNO = vomeronasal organ; VNC = vomeronasal cartilage.

## Discussion

### Comparative VNO morphology in lutrines and other mustelids

Overall, the evidence presented here suggests that the VNO is present in freshwater otters but lack some of the derived features that diagnose the ‘rudimentary’ VNO in harbor seals (Table 1). It differs from the VNO of the American mink in having a smaller lumen relative to the body of the organ, no discernible neurovasculature surrounding the VND lumen, and increased empty space surrounding the VNO within the cartilaginous capsule. The functional implications of the variation in VND lumen morphology and the expanded capsular space between the two mustelids and the river otter described here are not yet understood. Previous studies have noted that a relatively dilated VNO lumen may indicate a thinner epithelial layer and, therefore, reduced function (Smith et al., 2024). Based on the comparative diceCTs of *N. vison* and *L. canadensis*, the lumen of *L. canadensis* appears to be less dilated than that of *N. vison*. This may reflect the observation of the more ‘tubular’ or circular-shaped lumen, also seen in the ferret *M. furo*, rather than an actual difference in volume due to dilation (Weiler et al., 1999). The shape of the VNO lumen in the North American river otter *L. canadensis* was only discernible when image contrast was increased and was smaller than in the other two species. Its VNO lumen and/or actual VNO may be reduced in size or compressed, creating the empty space observed between the outside of the VNO and the VNG or VNC. This suggests that the VNO is reduced in the North American river otter but does not affect the surrounding cartilage capsule that contains it. While vasculature was, presumably, observable in the *N. vison* diceCT, no vasculature was identifiable in the *L. canadensis* or the *N. frenata* diceCT VNO.

Lutrines appeared to have shorter and broader VNG compared to other caniforms. The reduction in VNG length may be produced by covariation or constraint with certain aspects of skull morphology (e.g., relative snout length, extent of nasal turbinates). Kondoh et al. (2025) suggest that such structural constraints may have driven the shift in VNO position to be housed entirely within the incisive canal in Steller sea lions. However, the description of a defective septum resulting in unaffected VNO morphology in domestic dogs suggests that the organ may not strongly covary with specific regions or features within the nasal cavity (Dzięcioł et al., 2020).

### Structure-function and phenotype-genotype decoupling in the accessory olfactory system

VNG length variation may produce differences in relative VNO morphology and therefore, in its function. Shorter VNOs may have reduced surface area for sensory epithelia than relatively longer VNOs. However, this assumes a one-to-one relationship between morphology and function, which is often complicated by kinematics or behavior. For example, the flehmen response, wherein the lips are raised to bring air over the VNO, theoretically supports the function of the organ, but the presence of this behavior is not detectable by the morphology of the VNS overall. In caniforms, this behavior is largely absent, with the exception of the honey badger (*Mellivora capensis*), a fossorial mustelid (Begg et al., 2003) and reports in the polar bear (Stirling, 2016) (but see the pseudo-flehmen response observed in juvenile *E. lutris*, Island et al. 2017). It is also possible that, even with a thorough understanding of both VNS genotype and phenotype, neither may accurately predict the range of detectable odorants in mammal species (Chengetanai et al., 2020). Ultimately, several interacting mechanisms supporting chemosensory function (morphology, genes, behavior) make interpretations about relative function more complex.

Further evaluation of whether a shortened VNO observed in Lutrinae correlates with reduced function would require a more diverse sample that preserves the soft tissues of the VNO. However, the reduction of both VNG length and MOB volume in Lutrines (Gittleman, 1991) suggests a synchronous reduction of the two systems in the transition to an aquatic environment (Suárez et al., 2012). This is noteworthy given that the relationship among components of the VNS (i.e. genotype and phenotype, including morphological structures and behavioral outputs of the AOB and VNO) can evolve in a mosaic manner between and within the two systems (this study; Yohe & Kroll, 2018; Yohe et al., 2020; Smith et al., 2024). The observed reduction of both the MOS and AOS in Lutrinae suggests that, at this phylogenetic scale, phenotypic change associated with their ecological transition is sufficiently captured (Garrett & Steiper, 2014; Yohe et al., 2020).

The current literature does not present a strong consensus on the correlation between VNS phenotype and genotype: some studies support a strong correspondence (Garrett & Steiper 2014; Ibarra-Soria et al. 2014), where others demonstrate that the relationship weakens or becomes more complex at systemic or phylogenetic scales (Yohe et al., 2019; Yohe et al. 2020; Yohe & Krell 2023; Smith et al. 2024). Such correlations touch on the idea of phenogenetic drift (Weiss & Fullerton, 2000), which theorizes that selection differentially acts upon those two components. The concept of phenogenetic drift suggests that different genotypes produce the same phenotype, where phenotypes remain fixed or conserved while the potentially multi-loci genotypes underlying them change with time and space (Weiss & Fullerton, 2000). This typically applies when phenotypes are strongly selected for across different populations. The present study suggests the opposite dynamic in which pinnipeds and lutrines present with similar genotypes (reduced functional VNO genes, Yu et al. 2010; Zhao et al., 2020) and dissimilar phenotypes (Kondoh et al., 2024). Beyond their relevance to studies of the VNS, carnivorans, and secondarily aquatic mammals, these findings contribute to broader biological theory by supporting a mosaic relationship between chemosensory genotype and phenotype. At the taxonomic scale of Lutrinae within Caniformia, our results align with evidence that genotype-phenotype correspondence in chemosensory systems weakens at finer phylogenetic scales (Yohe et al., 2020; Yohe & Krell, 2023) and mirrors patterns of component-level VNS mosaicism described in bats (Smith et al., 2024).

A potential explanation for the longevity of VNS phenotypes following or during periods of genomic simplification is that they are, at least in part, patterned by surrounding tissues of the palate during embryological development. In this paradigm, information required for patterning the VNS is not only contained within genes expressed in the VNO or vomeronasal epithelia, but by the signaling environment of the developing palate as a whole. A growing body of literature suggests complex signaling dynamics exist between tissues that develop in close spatial proximity (e.g., Fabbri et al., 2017). Foster et al. (2024) identify a correlation wherein reptile lineages that undergo a radical restructuring of the palate also lose many, if not all, aspects of the VNS in adult phenotypes. Interestingly, embryos of said lineages actually exhibit transitory VNOs in early development, implying that the information required for their early patterning is present, but is disrupted by derived palatal morphogenesis. These insights have relevance for the retention of VNS phenotypes in mammalian lineages that have a reduced gene repertoire. Sampling of palatal development in these taxa, particularly those with morphologies related to an aquatic existence, might reveal explanatory factors for the presence or absence of VNS phenotypes.

### Hypothesized mechanisms for the derived VNS in pinnipeds

Generally, throughout the literature, a trait that appears particularly relevant to the function of the AOS is its connection to the oral cavity via the ID (Sanmartín-Vázquez et al., 2024). In dogs (*Canis familiaris*), the ID communicates between the two cavities even at birth (Sanmartín-Vázquez et al., 2024)., and one of the derived traits of the VNO in harbor seals (*Phoca vitulina*) is a lack of connection between the VNO and the oral cavity (although the incisive canal is still present; Kondoh et al., 2024). The retention of communication between the oral cavity and the nasal cavity where the VNO is housed in Steller sea lions (*Eumetopias jubatus*) provides increasing evidence that this feature serves as additional anatomical support for the presence of a patent VNO (Kondoh et al., 2025).

The loss of sensory-functional tissues in the derived VNO in harbor seals (*Phoca vitulina*) may be due to various selective, adaptive, or structural changes. One hypothesis is that the nasal cavity requires increased mucosal secretions to maintain a protective layer in salt water (Kondoh et al., 2024). This may be compounded by the loss of the nasolacrimal duct in pinnipeds (Berta et al., 2018; Colitz, 2022). However, this would not explain why phocids (such as the harbor seals) have derived VNOs while otariids do not, as all pinnipeds have lost the nasolacrimal duct and nearly all inhabit salt water. Another alternative explanation proposed is that otariids are more gregarious than phocids and generally spend more time on land - a combination of increased opportunities for social communication and other behavioral and biological factors creates selective pressure in maintaining a functional VNS (Kondoh et al., 2025).

Another alternative explanation for the transformation seen in pinnipeds reflects changes in their respiration. In horses, which are obligate nasal breathers, the connection between their oral and nasal cavities (ID) closes, although their VNO remains, and they retain a flehmen response (Mader, 2019). Pinnipeds, along with other marine mammals, highly prioritize nasal breathing, as the nose is superiorly located on the head relative to the oral cavity/mouth requiring them to raise less of their head from the water to inhale (Maust-Mohl et al., 2019; Reidenberg & Laitman, 2024). Kondoh et al. (2025) hypothesize that the derived VNO of the Steller sea lion (*Eumetopias jubatus*) functions through air intake via the oral cavity rather than relying on a venous pump. However, the extent to which pinnipeds are obligate nasal breathers remains unclear, and the literature lacks comprehensive reports on panting or mouth breathing across species, further complicating interpretations of VNO function in this group. In other aquatic mammals, such as cetaceans, evolutionary pressures have led to a significant reorganization of the facial skeleton, repositioning the nasal opening dorsally, which allows for efficient breathing at the water’s surface without full emergence (Berta et al., 2014; Maust-Mohl et al., 2019). This illustrates how obligate nasal breathing and associated craniofacial adaptations have evolved in response to the demands of an aquatic environment, further underscoring the need to examine whether similar respiratory constraints or strategies exist in pinnipeds.

Variation in the VNO may also correlate with changes to the role of respiration in thermoregulation. Pinnipeds appear to lack a panting mechanism for heat dissipation (Khamas et al., 2012), with the exception of the Northern Fur seal (*Callorhinus ursinus)* (Bartholomew & Wilke, 1956)—although future behavioral studies may find more exceptions to this observation. Some have suggested that control of the arterial supply in the nasal mucosa may be an alternative mechanism for regulating expired air temperature in pinnipeds (Folkow, 1992). Additionally, correlations have been found between the number of respiratory turbinates and climate within pinnipeds (Mason et al., 2020). Environmental temperature also differs across the range of phocids and otariids making an ecological correlation between the presence of a derived (phocid) and non-derived (otariid) VNS plausible. It has been documented that cold, dry air increases nasal epithelial shedding in humans (Cruz et al., 2006). The loss of any mouth-breathing and the potential for increased nasal mucosa to cope with colder temperatures, compounded with the loss of the nasolacrimal duct, may produce the selective pressures that have shaped the derived condition of the Harbor seal VNO. Future studies should also investigate whether AOS structure or function covaries with environmental factors or other aspects of the nasal cavity (e.g., width or turbinate area).

More broadly speaking, our understanding of some of the more specific functional mechanisms of the VNS is largely limited to studies in rodent models (Salazar & Sánchez Quinteiro, 2009). While these studies provide a great wealth of information regarding the VNO and AOS, it is evident that our limited knowledge affects assessments of the characteristics of the non-rodent mammalian AOS (Salazar & Sánchez Quinteiro, 2009). For example, the ferret VNO, which is smaller than that found in rodents, is described as rudimentary by Weiler and colleagues (1999), despite the lack of research on the phylogenetic structure and diversity of VNS structures in caniforms. These observations ultimately reflect an assumption that any regression of the AOS relative to the rat model may reflect a non-functional system. However, comparison within a phylogenetic context of caniforms highlights that their morphology reflects their own evolutionary specialization relative to rodents, some of which can be described as potential ‘regressions’ (Salazar et al., 2012). Future studies on the influence of ecology and phylogeny on the diversity of AOS structure and function in mammals require a broader integrative approach, examining the greater morphological diversity of this system within its phylogenetic context. This investigation of the VNS in secondarily aquatic caniforms is essential to understanding the response of mammalian sensory ecology and anatomy to ecological change.

### The definition of the ‘functional’ VNO

Future research is needed to standardize the terminology used to relate VNO structure to function. The literature currently employs a wide range of overlapping and sometimes ambiguous descriptors, including distinctions such as “functional” versus “non-functional” and “vestigial” or “rudimentary” versus “well-developed” or “true” VNOs (Smith et al., 2014).

Recent research has attempted to provide more impactful definitions of a functional VNO as “a neuroepithelium is essential to VNO function as a chemosensory organ, and we use the term ’neuroepithelial VNO’ to refer to those that possess both a neuroepithelium and bundle axons departing the basal aspect of the neuroepithelium. Based on our observations of the morphology, we might refer to bilateral epithelial tubes lacking a neuroepithelium and coextensive with VNCs as “putative rudimentary VNOs” (Smith et al., 2024), in contrast to the “true” VNS defined by Salazar and Sánchez-Quinteiro (2009) as comprising three components (the VNO, accessory olfactory bulb, and vomeronasal amygdala) with corresponding nerves and connections.

Patterns observed in bats highlight the complexity of the relationships between AOS anatomical components that may reflect ‘functional’ versus ‘rudimentary’ VNOs (Smith et al., 2024). Bats that lack AOBs and functional VNO genes retain neurological and vascular VNO structures that persist into adulthood. Bats without a ‘functional’ VNO (“possess(ing) both a neuroepithelium and bundle axons departing the basal aspect of the neuroepithelium”) often retain multiple structural elements of the VNO (Smith et al., 2024). This would mean that the definition of the ‘true’ VNO outlined by Salazar and Sánchez-Quinteiro (2009) would define those bats as likely having “true” VNOs, while Smith et al. (2024) would describe them as “putative rudimentary VNOs”. The presence of neurovascular tissue as a criterion for a functional VNO/AOS is complicated further by the cases of nerves from the VNO communicating with the MOB instead of the AOB in species that lack an AOB (Salazar & Sánchez-Quinteiro, 2009). This suggests that the presence of neural tissue and vascular structures may be a poor predictor of a “functional” VNO in carnivorans or, more generally, across Mammalia, although further research is needed. In general, future research should explicitly investigate the VNS across its various levels of organization (genotype and multiple aspects of phenotype) before defining groups as having “non-functional” VNS.

### Other future directions

Evaluating the soft-tissue morphology of the sea otter VNO would greatly contribute to further determining correlations between ecological, genetic, and anatomical transformations in the VNS. Sea otters (*Enhydra lutris*) are widely regarded as ecologically distinct from freshwater otters (Estes, 1989; Bird et al., 2020), spending significantly less time on land and exhibiting minimal reliance on terrestrial environments for feeding, reproduction, offspring care, or other ecological functions. In birds, gene patterns suggest that varying degrees and types of aquatic lifestyles are associated with corresponding, gradational changes in the olfactory gene repertoire (Lu et al., 2016), a pattern that may likewise be reflected in the AOS structures of aquatic mammals. Based on the derived VNG morphology compared to other otters and their lifestyle, sea otters would be the most likely lutrine to converge on the AOS morphology seen in the Harbor seal (*Phoca vitulina*). It is worth noting that captive juvenile sea otters have been observed exhibiting a flehmen-like response (Island et al., 2017); however, the relationship between this behavioral observation and the state of the sea otter VNS is not clearly understood.

Another future contribution that would significantly contribute to our understanding of the AOS in secondarily aquatic caniforms would be a thorough comparative description or identification of the AOB. While here we identify and describe soft tissue of the VNO in the North American river otter, there are no data in the literature describing their AOB. A description of AOB structures has previously only been achievable through histology or dissection (Ortiz-Leal et al., 2020, 2024). However, recent work (Gignac et al., 2021; Straight et al., 2024) presents the exciting possibility of identifying and describing it in detail using diceCT techniques.

In addition to otters, greater comparative sampling across phocid seals would be highly beneficial, as they represent immense ecological and taxonomic diversity that is underrepresented in the descriptive literature on VNS morphology. While describing or investigating these morphologies using diceCT or histology would be valuable, some aspects of the VNS may potentially be represented using osteological correlates. For example, it is hypothesized that the AOB may be observable on the endocast of some otariid pinnipeds (Loza et al., 2023), and the presence of a patent communication between the oral and nasal cavities can be investigated in hard tissues via the incisive canal.

While the VNG serves as a strong proxy for dimensions of the VNO in primates (Smith et al., 2011; Garrett et al., 2013; Garrett, 2015), it is possible, considering the condition seen in pinnipeds (Kondoh et al., 2024), that the VNG is simply an osteological correlate for the VNC in carnivorans. While this is still promising, as the VNC is strongly correlated with the presence of a VNO, or at least derived structures homologous to a VNO, it means that the presence of a VNG is not necessarily an osteological correlate of a *functional* VNO. The harbor seal (*Phoca vitulina*) retains what is regarded as a rudimentary (or exapted) VNO (secretory tissue) and VNC (Kondoh et al., 2024). Further, the presence of the VNG as evidence for a VNO in pinnipeds is complicated by the fact that the Steller sea lion (*Eumetopias jubatus*) VNO is present but is constrained to the incisive canal and does not extend into the nasal cavity (Kondoh et al., 2025). While this study confirms the presence of a VNO and establishes that the VNG is a strong correlate for the presence of VNO-derived or homologous tissue, further study is required to better reconstruct histo-morphological features that better capture ‘functional’ aspects of the VNO. These would include examination of the epithelium expressed throughout the VNO lumen and a detailed study of VNO neurovasculature.

## Conclusion

Here, we established the presence of a vomeronasal organ in one species, *Lontra canadensis,* a member of the family Lutrinae for which very little was known regarding the functional morphology of their accessory olfactory system. The correlation between a VNO and VNG across the sample of Caniformia suggests that the VNG can offer insight into the evolution of the AOS within carnivorans. Our findings suggest that there is phenogenetic drift within the caniform VNS, at least in secondarily aquatic species, highlighting the mosaic, irregular, and complex nature of structural and genetic change in the VNS. Future research should prioritize characterizing the histo-morphological aspects of the VNS in secondarily aquatic carnivorans to elucidate phenogenetic patterns, especially during ecological transitions. Such work would also facilitate comparisons with VNS changes documented in other mammalian clades (Yohe & Krell, 2023) and may help identify generalized characteristics of the system across mammals or even tetrapods.

## Acknowledgements

We thank Drs. Gabriel Bever, Alistair Evans, and James Rule for reviewing and discussing early editions of this manuscript as part of S. M. Palmer’s dissertation completed at Johns Hopkins University. We also acknowledge the Johns Hopkins University Materials Characterization and Processing (MCP) facility for access to equipment and technical support. We thank both Arianna Harrington (Southern Utah University) and Sandrine Ladevèze (MNHN Paris) for scanning and contributing specimens used in this study, as well as the numerous contributors to MorphoSource for access to other specimens (see Supplemental Table 1 for detailed MorphoSource acknowledgements).

## Sources cited

Allouch, G. M., & Alshanbari, F. A. (2024). Comparative Anatomy of the Vomeronasal Organ (VNO) in Sheep (Ovis aries) and Dogs (Canis familiaris) with Simple Reference to its Histological Structure and Vasculature Supply. International Journal of Morphology, 42(2).

Aron, C. (1979). Mechanisms of control of the reproductive function by olfactory stimuli in female mammals. Physiological Reviews, 59(2), 229–284.

Bartholomew, G. A., & Wilke, F. (1956). Body temperature in the northern fur seal, Callorhinus ursinus. Journal of Mammalogy, 37(3), 327–337.

Baum, M. J. (2012). Contribution of pheromones processed by the main olfactory system to mate recognition in female mammals. Frontiers in neuroanatomy, 6, 20.

Baum, M. J., & Cherry, J. A. (2015). Processing by the main olfactory system of chemosignals that facilitate mammalian reproduction. Hormones and behavior, 68, 53–64.

Baum, M. J., & Kelliher, K. R. (2009). Complementary roles of the main and accessory olfactory systems in mammalian mate recognition. Annual Review of Physiology, 71(1), 141–160.

Begg, C. M., Begg, K. S., Du Toit, J. T., & Mills, M. G. L. (2003). Scent-marking behaviour of the honey badger, Mellivora capensis (Mustelidae), in the southern Kalahari. Animal behaviour, 66(5), 917–929.

Bendel, E. M., Kammerer, C. F., Kardjilov, N., Fernandez, V., & Fröbisch, J. (2018). Cranial anatomy of the gorgonopsian Cynariops robustus based on CT-reconstruction. PLoS One, 13(11), e0207367.

Berta, A., Churchill, M., & Boessenecker, R. W. (2018). The Origin and Evolutionary Biology of Pinnipeds. Annual Review of Earth and Planetary Sciences, 46, 203–228.

Berta, A., Ekdale, E. G., & Cranford, T. W. (2014). Review of the cetacean nose: form, function, and evolution. The Anatomical Record, 297(11), 2205–2215.

Berta, A., Sumich, J. L., & Kovacs, K. M. (2014). Marine Mammals: Evolutionary Biology (3rd ed.). Academic Press.

Bird, D. J., Hamid, I., Fox-Rosales, L., & Van Valkenburgh, B. (2020). Olfaction at depth: Cribriform plate size declines with dive depth and duration in aquatic arctoid carnivorans. Ecology and Evolution, 10(14), 6929–6953.

Bird, D. J., Murphy, W. J., Fox-Rosales, L., Hamid, I., Eagle, R. A., & Van Valkenburgh, B. (2018). Olfaction written in bone: cribriform plate size parallels olfactory receptor gene repertoires in Mammalia. Proceedings of the Royal Society B: Biological Sciences, 285(1874), 20180100.

Caianiello, S. (2024). The Strange Story of Mosaic Evolution. In: Delisle, R.G., Esposito, M., Ceccarelli, D. (eds) Unity and Disunity in Evolutionary Biology. Springer, Cham. 10.1007/978-3-031-42629-2_13

Camilieri-Asch, V., Shaw, J. A., Mehnert, A., Yopak, K. E., Partridge, J. C., & Collin, S. P. (2020). diceCT: A valuable technique to study the nervous system of fish. Eneuro, 7(4).

Chengetanai, S., Bhagwandin, A., Bertelsen, M. F., Hård, T., Hof, P. R., Spocter, M. A., & Manger, P. R. (2020). The brain of the African wild dog. II. The olfactory system. Journal of Comparative Neurology, 528(18), 3285–3304.

Colitz, C. (2022). Ophthalmology of pinnipedimorpha: seals, sea lions, and walruses. In Wild and Exotic Animal Ophthalmology: Volume 2: Mammals (pp. 269–309). Cham: Springer International Publishing.

Collin, S. P., Yopak, K. E., Crowe-Riddell, J. M., Camilieri-Asch, V., Kerr, C. C., Robins, H., … & Chapuis, L. (2024). Bioimaging of sense organs and the central nervous system in extant fishes and reptiles in situ: A review. The Anatomical Record.

Crompton, A. W., Owerkowicz, T., Bhullar, B. A., & Musinsky, C. (2017). Structure of the nasal region of non-mammalian cynodonts and mammaliaforms: speculations on the evolution of mammalian endothermy. Journal of Vertebrate Paleontology, 37(1), e1269116.

Cruz, A. A., Naclerio, R. M., Proud, D., & Togias, A. (2006). Epithelial shedding is associated with nasal reactions to cold, dry air. Journal of allergy and clinical immunology, 117(6), 1351–1358.

Dawood, Y., Hagoort, J., Siadari, B. A., Ruijter, J. M., Gunst, Q. D., Lobe, N. H. J., … & Van Den Hoff, M. J. B. (2021). Reducing soft-tissue shrinkage artefacts caused by staining with Lugol’s solution. Scientific reports, 11(1), 19781.

de Ferran, V., Figueiro, H. V., de Jesus Trindade, F., Smith, O., Sinding, M. H. S., Trinca, C. S., … & Eizirik, E. (2022). Phylogenomics of the world’s otters. Current Biology, 32(16), 3650–3658.

De Vreese, S., Orekhova, K., Morell, M., Gerussi, T., & Graïc, J. M. (2023). Neuroanatomy of the cetacean sensory systems. Animals, 14(1), 66.

Debey, L. B., & Pyenson, N. D. (2013). Osteological correlates and phylogenetic analysis of deep diving in living and extinct pinnipeds: what good are big eyes?. Marine Mammal Science, 29(1), 48–83.

Dzięcioł, M., Podgórski, P., Stańczyk, E., Szumny, A., Woszczyło, M., Pieczewska, B., … & Wrzosek, M. A. (2020). MRI features of the vomeronasal organ in dogs (Canis familiaris). Frontiers in Veterinary Science, 7, 159.

Estes, J. A. (1989). Adaptations for aquatic living by carnivores. In Carnivore behavior, ecology, and evolution (pp. 242–282). Boston, MA: Springer US.

Estes, R. D. (1972). The role of the vomeronasal organ in mammalian reproduction. Evans, H. E. (1993). Miller’s Anatomy of the Dog (3rd ed.). W.B. Saunders.

Fabbri, M., Mongiardino Koch, N., Pritchard, A. C., Hanson, M., Hoffman, E., Bever, G. S., Balanoff, A. M., Morris, Z. S., Field, D. J., Camacho, J. and Rowe, T. B. 2017. The skull roof tracks the brain during the evolution and development of reptiles including birds. Nature ecology & evolution, 1(10), 1543–1550.

Folkow, L. P. (1992). Adrenergic vasomotor responses in nasal mucosa of hooded seals. American Journal of Physiology-Regulatory, Integrative and Comparative Physiology, 263(6), R1291–R1297.

Foster, W., Gensbigler, P., Wilson, J. D., Smith, RMH., Lyson, T. R., & Bever, G. S. 2024. Cranial Anatomy of the Triassic Rhynchosaur *Mesosuchus browni* based on computed tomography, with a discussion of the vomeronasal system and its deep history in Reptilia. Zoological Journal of the Linnean Society, 201(4), zlae097.

Garrett, E. C. (2015). Was there a sensory trade-off in primate evolution? The vomeronasal groove as a means of understanding the vomeronasal system in the fossil record. City University of New York.

Garrett, E. C., Dennis, J. C., Bhatnagar, K. P., Durham, E. L., Burrows, A. M., Bonar, C. J., … & Smith, T. D. (2013). The vomeronasal complex of nocturnal strepsirhines and implications for the ancestral condition in primates. The Anatomical Record, 296(12), 1881–1894.

Gignac, P. M., Kley, N. J., Clarke, J. A., Colbert, M. W., Morhardt, A. C., Cerio, D., … & Witmer, L. M. (2016). Diffusible iodine-based contrast-enhanced computed tomography (diceCT): an emerging tool for rapid, high-resolution, 3-D imaging of metazoan soft tissues. Journal of anatomy, 228(6), 889–909.

Gignac, P. M., O’Brien, H. D., Sanchez, J., & Vazquez-Sanroman, D. (2021). Multiscale imaging of the rat brain using an integrated diceCT and histology workflow. Brain Structure and Function, 226(7), 2153–2168.

Gittleman, J. L. (1991). Carnivore olfactory bulb size: allometry, phylogeny and ecology. Journal of Zoology, 225(2), 253–272.

Hadden, P. W., & Zhang, J. (2023). An overview of the penguin visual system. Vision, 7(1), 6.

Hecker, N., Lächele, U., Stuckas, H., Giere, P., & Hiller, M. (2019). Convergent vomeronasal system reduction in mammals coincides with convergent losses of calcium signalling and odorant-degrading genes. Molecular Ecology, 28(16), 3656–3668.

Hillenius, W. J. (2000). Septomaxilla of nonmammalian synapsids: soft-tissue correlates and a new functional interpretation. Journal of morphology, 245(1), 29–50.

Hughes, N. K., Price, C. J., & Banks, P. B. (2010). Predators are attracted to the olfactory signals of prey. PLoS One, 5(9), e13114.

Island, H. D., Wengeler, J., & Claussenius-Kalman, H. (2017). The flehmen response and pseudosuckling in a captive, juvenile Southern sea otter (Enhydra lutris nereis). Zoo Biology, 36(1), 30–39.

Kelliher, K. R., Baum, M. J., & Meredith, M. (2001). The ferret’s vomeronasal organ and accessory olfactory bulb: effect of hormone manipulation in adult males and females. The Anatomical Record: An Official Publication of the American Association of Anatomists, 263(3), 280–288.

Keverne, E. B. (2004). Importance of olfactory and vomeronasal systems for male sexual function. Physiology & behavior, 83(2), 177–187.

Khamas, W. A., Smodlaka, H., Leach-Robinson, J., & Palmer, L. (2012). Skin histology and its role in heat dissipation in three pinniped species. Acta Veterinaria Scandinavica, 54, 1–10.

Kondoh, D., Tonomori, W., Iwasaki, R., Tomiyasu, J., Kaneoya, Y., Kawai, Y. K., … & Kobayashi, M. (2024). The vomeronasal organ and incisive duct of harbor seals are modified to secrete acidic mucus into the nasal cavity. Scientific Reports, 14(1), 11779.

Kondoh, D., Tonomori, W., Iwasaki, R., Tomiyasu, J., Kaneoya, Y., Li, H., … & Kobayashi, M. (2025). The vomeronasal system of the Steller sea lion. Journal of Anatomy.

Leng, L., & Shi, L. (2025). How Foraging Mode Sculpts Sensory Systems: Morphological Evidence From DiceCT and Histology in Sympatric Lizards. Ecology and Evolution, 15(8), e72042.

Liu, A., He, F., Shen, L., Liu, R., Wang, Z., & Zhou, J. (2019). Convergent degeneration of olfactory receptor gene repertoires in marine mammals. BMC genomics, 20, 1–14.

Loza, C. M., Sánchez-Villagra, M. R., Scarano, A. C., Romero, M., Barbeito, C. G., & Carlini, A. A. (2023). The brain of fur seals, seals, and walrus (Pinnipedia): A comparative anatomical and phylogenetic study of cranial endocasts of semiaquatic mammals. Journal of Mammalian Evolution, 30(4), 1011–1028.

Lu, Q., Wang, K., Lei, F., Yu, D., & Zhao, H. (2016). Penguins reduced olfactory receptor genes common to other waterbirds. Scientific Reports, 6(1), 31671.

Mader, B. J. (2019). The narial morphology of Metarhinus and Sphenocoelus (Mammalia, Perissodactyla, Brontotheriidae).

Mahdy, E. A., & Mohamed, S. K. A. (2019). Comparative morpho-histological analysis on the vomeronasal organ and the accessory olfactory bulb in Balady dogs (Canis familiaris) and New Zealand rabbits (Oryctolagus cuniculus). Journal of Advanced Veterinary and Animal Research, 6(4), 506.

Mason, M. J., Wenger, L. M., Hammer, Ø., & Blix, A. S. (2020). Structure and function of respiratory turbinates in phocid seals. Polar Biology, 43, 157–173.

Niimura, Y., & Nei, M. (2007). Extensive gains and losses of olfactory receptor genes in mammalian evolution. PloS one, 2(8), e708.

Ortiz-Leal, I., Torres, M. V., Barreiro-Vázquez, J. D., López-Beceiro, A., Fidalgo, L., Shin, T., & Sanchez-Quinteiro, P. (2024). The vomeronasal system of the wolf (Canis lupus signatus): The singularities of a wild canid. Journal of Anatomy, 245(1), 109–136.

Ortiz-Leal, I., Torres, M. V., Villamayor, P. R., Fidalgo, L. E., López-Beceiro, A., & Sanchez-Quinteiro, P. (2022). Can domestication shape Canidae brain morphology? The accessory olfactory bulb of the red fox as a case in point. Annals of Anatomy-Anatomischer Anzeiger, 240, 151881.

Ortiz-Leal, I., Torres, M. V., Villamayor, P. R., López-Beceiro, A., & Sanchez-Quinteiro, P. (2020). The vomeronasal organ of wild canids: the fox (Vulpes vulpes) as a model. Journal of Anatomy, 237(5), 890–906.

Paulina-Carabajal, A., Acosta-Hospitaleche, C., & Yury-Yáñez, R. E. (2015). Endocranial morphology of Pygoscelis calderensis (Aves, Spheniscidae) from the Neogene of Chile and remarks on brain morphology in modern Pygoscelis. Historical Biology, 27(5), 571–582.

Pihlström, H., Thewissen, J. G. M., & Nummela, S. (2008). Comparative anatomy and physiology of chemical senses in aquatic mammals. Sensory evolution on the threshold: Adaptations in secondarily aquatic vertebrates, 95–109.

Poo, C., Agarwal, G., Bonacchi, N., & Mainen, Z. F. (2022). Spatial maps in piriform cortex during olfactory navigation. Nature, 601(7894), 595–599.

Reidenberg, J. S., & Laitman, J. T. (2025). Review of respiratory anatomy adaptations in whales. The Anatomical Record, 308(4), 1179–1213.

Rolfe, S., Pieper, S., Porto, A., Diamond, K., Winchester, J., Shan, S., … & Maga, A. M. (2021). SlicerMorph: An open and extensible platform to retrieve, visualize and analyse 3D morphology. Methods in Ecology and Evolution, 12(10), 1816–1825.

Salazar, I., & Sánchez Quinteiro, P. (2009). The risk of extrapolation in neuroanatomy: the case of the mammalian vomeronasal system. Frontiers in neuroanatomy, 3, 982.

Salazar, I., Cifuentes, J. M., & Sánchez-Quinteiro, P. (2013). Morphological and immunohistochemical features of the vomeronasal system in dogs. The Anatomical Record: Advances in Integrative Anatomy and Evolutionary Biology, 296(1), 146–155.

Salazar, I., Lombardero, M., Cifuentes, J. M., Quinteiro, P. S., & Alemañ, N. (2003). Morphogenesis and growth of the soft tissue and cartilage of the vomeronasal organ in pigs. Journal of Anatomy, 202(6), 503–514.

Salazar, I., Quinteiro, P. S., & Cifuentes, J. M. (1995). Comparative anatomy of the vomeronasal cartilage in mammals: mink, cat, dog, pig, cow and horse. Annals of Anatomy-Anatomischer Anzeiger, 177(5), 475–481.

Sanmartín-Vázquez, E., Ortiz-Leal, I., Torres, M. V., Kalak, P., Kubiak-Nowak, D., Dzięcioł, M., & Sanchez-Quinteiro, P. (2024). Functional Role of the Incisive Duct in Neonatal Dogs. Cells Tissues Organs, 1–30.

Smith, T. D., Corbin, H. M., King, S. E., Bhatnagar, K. P., & DeLeon, V. B. (2021). A comparison of diceCT and histology for determination of nasal epithelial type. PeerJ, 9, e12261. the language of anatomical reduction. *The Anatomical Record*, *297*(11), 2196-2204.

Smith, T. D., Downing, S. E., Rosenberger, V. B., Loeffler, J. R., King, N. A., Curtis, A. A., … & Santana, S. E. (2024). Functional microanatomy of the vomeronasal complex of bats. The Anatomical Record.

Smith, T. D., Garrett, E. C., Bhatnagar, K. P., Bonar, C. J., Bruening, A. E., Dennis, J. C., … & Morrison, E. E. (2011). The vomeronasal organ of New World monkeys (Platyrrhini). The Anatomical Record: Advances in Integrative Anatomy and Evolutionary Biology, 294(12), 2158–2178.

Smith, T. D., Laitman, J. T., & Bhatnagar, K. P. (2014). The shrinking anthropoid nose, the human vomeronasal organ, and the language of anatomical reduction. The Anatomical Record, 297(11), 2196–2204.

Stirling, I., Spencer, C., & Andriashek, D. (2016). Behavior and activity budgets of wild breeding polar bears (Ursus maritimus). Marine Mammal Science, 32(1), 13–37.

Straight, P. J., Gignac, P. M., & Kuenzel, W. J. (2024). A histological and diceCT-derived 3D reconstruction of the avian visual thalamofugal pathway. Scientific Reports, 14(1), 8447.

Suárez, R., García-González, D., & De Castro, F. (2012). Mutual influences between the main olfactory and vomeronasal systems in development and evolution. Frontiers in neuroanatomy, 6, 50.

Switzer, III, R. C., Johnson, J. I., & Kirsch, J. A. (1980). Phylogeny through brain traits: relation of lateral olfactory tract fibers to the accessory olfactory formation as a palimpsest of mammalian descent. Brain, Behavior and Evolution, 17(5), 339–363.

Tirindelli, R. (2021). Coding of pheromones by vomeronasal receptors. Cell and Tissue Research, 383(1), 367–386.

Tomiyasu, J., Kondoh, D., Sakamoto, H., Matsumoto, N., Sasaki, M., Kitamura, N., … & Matsui, M. (2017). Morphological and histological features of the vomeronasal organ in the brown bear. Journal of anatomy, 231(5), 749–757.

Waku, D., Segawa, T., Yonezawa, T., Akiyoshi, A., Ishige, T., Ueda, M., … & Sasaki, T. (2016). Evaluating the phylogenetic status of the extinct Japanese otter on the basis of mitochondrial genome analysis. PLoS One, 11(3), e0149341.

Weiler, E., Apfelbach, R., & Farbman, A. I. (1999). The vomeronasal organ of the male ferret. Chemical senses, 24(2), 127–136.

Weiss, K. M., & Fullerton, S. M. (2000). Phenogenetic drift and the evolution of genotype–phenotype relationships. Theoretical population biology, 57(3), 187–195.

Witmer, L. M., & Thomason, J. J. (1995). The extant phylogenetic bracket and the importance of reconstructing soft tissues in fossils. Functional morphology in vertebrate paleontology, 1, 19–33.

Yohe, L. R., & Krell, N. T. (2023). An updated synthesis of and outstanding questions in the olfactory and vomeronasal systems in bats: Genetics asks questions only anatomy can answer. The Anatomical Record, 306(11), 2765–2780.

Yohe, L. R., Fabbri, M., Hanson, M., & Bhullar, B. A. S. (2020). Olfactory receptor gene evolution is unusually rapid across Tetrapoda and outpaces chemosensory phenotypic change. Current Zoology, 66(5), 505–514.

Yohe, L. R., Hoffmann, S., & Curtis, A. (2018). Vomeronasal and olfactory structures in bats revealed by DiceCT clarify genetic evidence of function. Frontiers in neuroanatomy, 12, 32.

Yu, L., Jin, W., Wang, J. X., Zhang, X., Chen, M. M., Zhu, Z. H., … & Zhang, Y. P. (2010). Characterization of TRPC2, an essential genetic component of VNS chemoreception, provides insights into the evolution of pheromonal olfaction in secondary-adapted marine mammals. Molecular biology and evolution, 27(7), 1467–1477.

Zellmer, N. T., Timm-Davis, L. L., & Davis, R. W. (2021). Sea otter behavior: morphologic, physiologic, and sensory adaptations. In Ethology and behavioral ecology of sea otters and polar bears (pp. 23–55). Cham: Springer International Publishing.

Zhang, Z., & Nikaido, M. (2020). Inactivation of ancV1R as a predictive signature for the loss of vomeronasal system in mammals. Genome Biology and Evolution, 12(6), 766–778.

Zhao, H., Xu, D., Zhang, S., & Zhang, J. (2011). Widespread losses of vomeronasal signal transduction in bats. Molecular biology and evolution, 28(1), 7–12.

Zufall, F., Kelliher, K. R., & Leinders-Zufall, T. (2002). Pheromone detection by mammalian vomeronasal neurons. Microscopy research and technique, 58(3), 251–260.

