## Supplementary material for "Soft tissue morphology of the vomeronasal organ in *Lontra canadensis* and its osteological correlate: Implications for the evolution of the caniform accessory olfactory system": Suppl. Tables

Supplemental Table 1. Specimens used in this study.

| Specimen Number | Species |
|---|---|
| AMNH 89030 | *Ailuropoda melanoleuca* |
| AMNH M 35387 | *Ailurus fulgens* |
| RBINS 7838 | *Aonyx capensis* |
| MNHN ZM 1877-704 | *Arctonyx collaris* |
| YPM 007471 | *Canis familiaris (saluki)* |
| AMNH 80291 | *Canis lupus* |
| MNHN ZM 2000-646 | *Eira barbara* |
| MNHN ZM MO 1-35-124;<br>MNHN ZM MO 1962-1647 | *Enhydra lutris* |
| MNHN ZM 2001-1971 | *Galictis vittata* |
| AMNH 37433 | *Gulo gulo* |
| AMNH 35364 | *Helarctos malayanus* |
| AMNH 848807 | *Hydrictis maculicollis* |
| NCSM 19662 | *Lontra canadensis* |
| AMNH 48193 | *Lontra felina* |
| AMNH 98589 | *Lontra longicaudis* |
| MNHN ZM 2005-597 | *Lutra lutra* |
| USNM 259466 | *Lutra lutra chinensis* |
| MNHN ZM 1962-1646 | *Lutrogale perspicillata* |
| MNHN ZM 1962-1617 | *Lyncodon patagonicus* |
| NCSM 8313 | *Martes americana* |

| | |
|---|---|
| USNM 173295 | *Martes foina* |
| AMNH M 70603 | *Meles meles* |
| MNHN 1995-3150 | *Mellivora capensis* |
| MNHN 1929-376 | *Melogale moschata* |
| USNM 147523 | *Mephitis mephitis* |
| NCSM 16318 | *Mustela erminea* |
| NMS E190 | *Mustela lutreola* |
| NCSM 8102 | *Mustela nivalis* |
| AMNH 243106 | *Mustela putorius* |
| DUU-RFK-BA 049 | *Nasua narica* |
| UF 26043 | *Neogale frenata* |
| L-SPL1 | *Neovison vison* |
| USNM 254641 | *Nyctereutes procyonoides* |
| NCSM 5201 | *Pekania pennanti* |
| MNHN ZM 1943-146 | *Poecilictis libyca* |
| MNHN 1981-1371 | *Poecilogale albinucha* |
| USNM 337630 | *Potos flavus* |
| PSU ANTH 301.4 | *Procyon lotor* |
| USNM 304663 | *Pteronura brasiliensis* |
| UAM 88-50-364 | *Spilogale putorius* |
| NCSM 12989 | *Taxidea taxus* |
| AMNH 99308 | *Tremarctos ornatus* |
| USNM 213397 | *Ursus americanus* |

| | |
|---|---|
| USNM 82003 | *Ursus arctos* |
| ISM-mammals 001-05 | *Ursus maritimus* |
| UCLA 13112 | *Vulpes vulpes* |

Supplemental Table 2. DiceCT Specimens used:

| Specimen Number | Species |
|---|---|
| l-cet:001 | *Lontra canadensis* |
| YPM:VZ:YPM MAM 014733 | *Neogale frenata* |
| *None – Morphosource ID Media 000806888* | *Neogale vison* |

Other MorphoSource citations:

Some specimens originally appearing in Encephalic Arterial Canals and their Functional Significance and Scaling of bony canals for encephalic vessels in euarchontans: Implications for the role of the vertebral artery and brain metabolism, the collection of which was funded by NSF BCS 1825129 (to D.M. Boyer and A.R. Harrington).

Julian Avery and Timothy Ryan provided access to these data. The files were downloaded from www.MorphoSource.org, Duke University.

*Lontra canadensis* diceCT from Caitlin Yoakum, funded by NSF DDRIG 1944642.

*Neogale frenata* diceCT: Blackburn, D.C., et al. 2024. Increasing the impact of vertebrate scientific collections through 3D-imaging: the openVertebrate (oVert) Thematic Collections Network. BioScience 74: 169–186. https://doi.org/10.1093/biosci/biad120, funded via NSF DBI-1701714 ("oVert: TCN" parent grant).

Blaire Van Valkenburgh and Tim Rowe provided access to these data, with data collection funded by NSF IOB-0517748 and data upload to MorphoSource funded by DBI-1902242. The files were downloaded from www.MorphoSource.org, Duke University.
